# Locomotion-dependent use of geometric and body cues in humans mapping 3D space

**DOI:** 10.1101/2025.03.11.642663

**Authors:** Volker Reisner, Theo AJ Schäfer, Leonard König, Misun Kim, Christian F Doeller

**Affiliations:** Max Planck Institute for Human Cognitive and Brain Sciences, Leipzig, Germany; Institute of Psychology, University of Hamburg, Hamburg, Germany; Institute of Cognitive Neuroscience, University College London, London, United Kingdom; Kavli Institute for Systems Neuroscience, Centre for Neural Computation, The Egil and Pauline Braathen and Fred Kavli Centre for Cortical Microcircuits, Jebsen Centre for Alzheimer’s Disease, Norwegian University of Science and Technology, Trondheim, Norway

**Keywords:** cognitive map, spatial memory, 3D space, environmental geometry, locomotion, virtual reality, computational modeling

## Abstract

The ability to represent locations across multiple dimensions of space is a core function of cognitive maps. While the influence of boundary-dependent environmental geometry on spatial representations has been extensively studied in 2D spaces, less is known about the role of boundaries for volumetric spatial memory. Research in humans and other animals has demonstrated distinct processing of the vertical and horizontal spatial dimensions, likely related to species-specific modes of locomotion. Here, we investigate whether different locomotion modes, flying and walking, affect the use of vertical boundaries, leading to possibly distinct volumetric representations. In a Virtual Reality experiment, human participants memorized objects within a symmetric 3D enclosure, and then were asked to replace them in either the familiar or geometrically deformed environments. We found that the flying group exhibited lower vertical than horizontal spatial memory precision, whereas the walking group showed the opposite pattern, an effect related to using their body axis as a vertical “ruler”. Within deformed environments, object replacements in the flying group followed the predictions from a 3D-extended boundary-vector-cell-like computational model of spatial mapping that treated all boundaries equally, whereas those in the walking condition favored a modified model that prioritized the ground boundary. Our findings suggest that gravity-related movement constraints promote different utilization of geometric and body-related cues, resulting in flexible representations of volumetric space.

## Introduction

Representing locations in space constitutes a key function of the cognitive map, a mental model of the environment that supports flexible navigation (O’Keefe & Nadel, 1978; Tolman, 1948). Extensive research across species has demonstrated that cognitive maps are significantly influenced by environmental boundaries. These serve as highly stable reference cues, which separate space into navigable regions and define their geometric shape (Barry et al., 2007; Bellmund et al., 2019; Carpenter et al., 2015; Cheng, 1986; Hartley et al., 2004; Keinath et al., 2018, 2021; Krupic et al., 2015; Lee, 2017; O’Keefe & Burgess, 1996; Stensola et al., 2012). For instance, when the geometry of a familiar enclosure was changed from a square to a rectangle, both the firing patterns of rodent place cells and human spatial memory responses adapted according to the geometric properties of the new environment (O’Keefe & Burgess, 1996; Hartley et al., 2004). In the brain, these properties are signaled by boundary-vector cells (BVCs), which fire at specific allocentric direction and distance from the walls (Lever et al., 2009; Stewart et al., 2014). Notably, computational models mimicking the population activity of BVCs have been successful in predicting both rodent place cell activity and human spatial memory in geometrically deformed environments (Barry et al., 2006; De Cothi & Barry, 2020; Hartley et al., 2000; Hartley et al., 2004; O’Keefe & Burgess, 1996), suggesting a common neural mechanism for anchoring cognitive maps across species (Lee et al., 2018; Stangl et al., 2021).

How boundary information is utilized and encoded during navigation has mostly been studied on horizontal surfaces. However, while animals routinely traverse three-dimensional environments, little is known about how vertical boundaries, such as floors and ceilings, contribute to the formation of three-dimensional cognitive maps. Understanding the coding principles of the third spatial dimension is crucial for expanding our knowledge of how multi-dimensional cognitive maps support broader aspects of cognition, extending beyond physical navigation (Behrens et al., 2018; Bellmund et al., 2018).

The vertical spatial dimension is unique due to the influence of gravity. Flying animals can more readily defy gravity and approach boundaries along both horizontal and vertical dimensions to estimate distance. Place cell recordings in freely flying bats exhibited isotropic firing fields with similar resolution across spatial dimensions (Yartsev & Ulanovsky, 2013). In contrast, surface-dwelling animals are more constrained in their movements towards boundaries in the opposite direction of gravity (ceiling) and may need to rely more on vision for vertical distance estimation. However, depth perception along the vertical axis can be less accurate due to head tilt (Dhaliwal et al. 2002, Skerswetat et al. 2020), which may contribute to anisotropic spatial representations observed in both rats (Grieves et al., 2020; Hayman et al., 2011) and humans (Zwergal et al., 2016; Du & Mou, 2022; Hinterecker et al., 2018a; for diverging findings see Kim & Maguire, 2017; Hinterecker et al., 2018b). Given these cross-species differences, it was hypothesized that surface-dwelling animals such as ourselves, might have evolved to prioritize horizontal over vertical spatial information due to their natural mode of locomotion (Davis, Holbrook & de Perera, 2018).

Nonetheless, gravity can also serve as a useful cue for representing vertical space relative to the ground. This becomes particularly clear in environments with reduced gravity, like on space stations, where astronauts often report severe disorientation (Oman, 2007). On earth, gravity ensures stable sensory feedback from the vestibular system which, in primates, can signal tilt along the vertical dimension and is not available for horizontal directions (Angelaki & Cullen, 2008). At the same time, gravity consistently defines a stable vertical axis which is aligned with the upright human body axis that rises from the ground. When moving on a horizontal plane, the body’s orthogonal relationship to the ground remains constant, providing a reliable point of reference and, thus, potentially making it easier to estimate height between the ground and one’s own body dimensions (e.g. the head).

In the present study, we sought to understand the role of environmental boundaries in the representation of three-dimensional space, considering two modes of locomotion: One that allows for 3D navigation similar to winged animals (flying group) and one that is restricted to the ground reflecting naturalistic human locomotion (walking group). This allowed us to investigate how humans integrate boundary- and body-related cues under different degrees of freedom for moving against gravity. Furthermore, we deformed the geometry of the environment to investigate the causal influence of boundaries on spatial memory. By applying computational geometric models (Hartley et al., 2004) extended into 3D space, we uncovered distinct strategies depending on the mode of locomotion: those who walked tended to rely more on ground- and body-based references for vertical estimation, while those who flew exhibited more equal encoding relative to boundaries across dimensions.

## Results

We analyzed data from 77 healthy young participants (40 female, 36 male, 1 non-binary; Age: 26.2 ± 4.5 years, 19-35 years) performing a 3D object-location memory task in an immersive Virtual Reality (VR) with Motion Capture (MoCap) technology (Fig. 1A-B). During training, participants first learned 3D locations of 6 free-floating objects within a cubic baseline environment. This manipulation is novel compared to previous studies where objects were lying on the walls, floor, or discretized 3D structure (Kim & Maguire, 2017; Zwergal et al., 2016; Hinterecker, 2018b; Du & Mou, 2022). During the subsequent test, participants were asked to replace the objects from pseudo-random start locations. We probed their volumetric spatial memory in either the familiar or geometrically deformed environments (Fig. 1C). While for some participants the deformed environment was stretched (groups 1 & 2) and for others it was compressed (groups 3 & 4), all participants encountered deformations along both the horizontal and vertical dimensions. Ultimately, participants completed the experiment in one of two locomotion modes, with walking participants (groups 1 & 3) performing natural human locomotion and virtual flying participants mimicking natural locomotion of winged animals (groups 2 & 4; Fig. 1D). The full experimental design is shown in Fig 1C.

**Figure 1.**
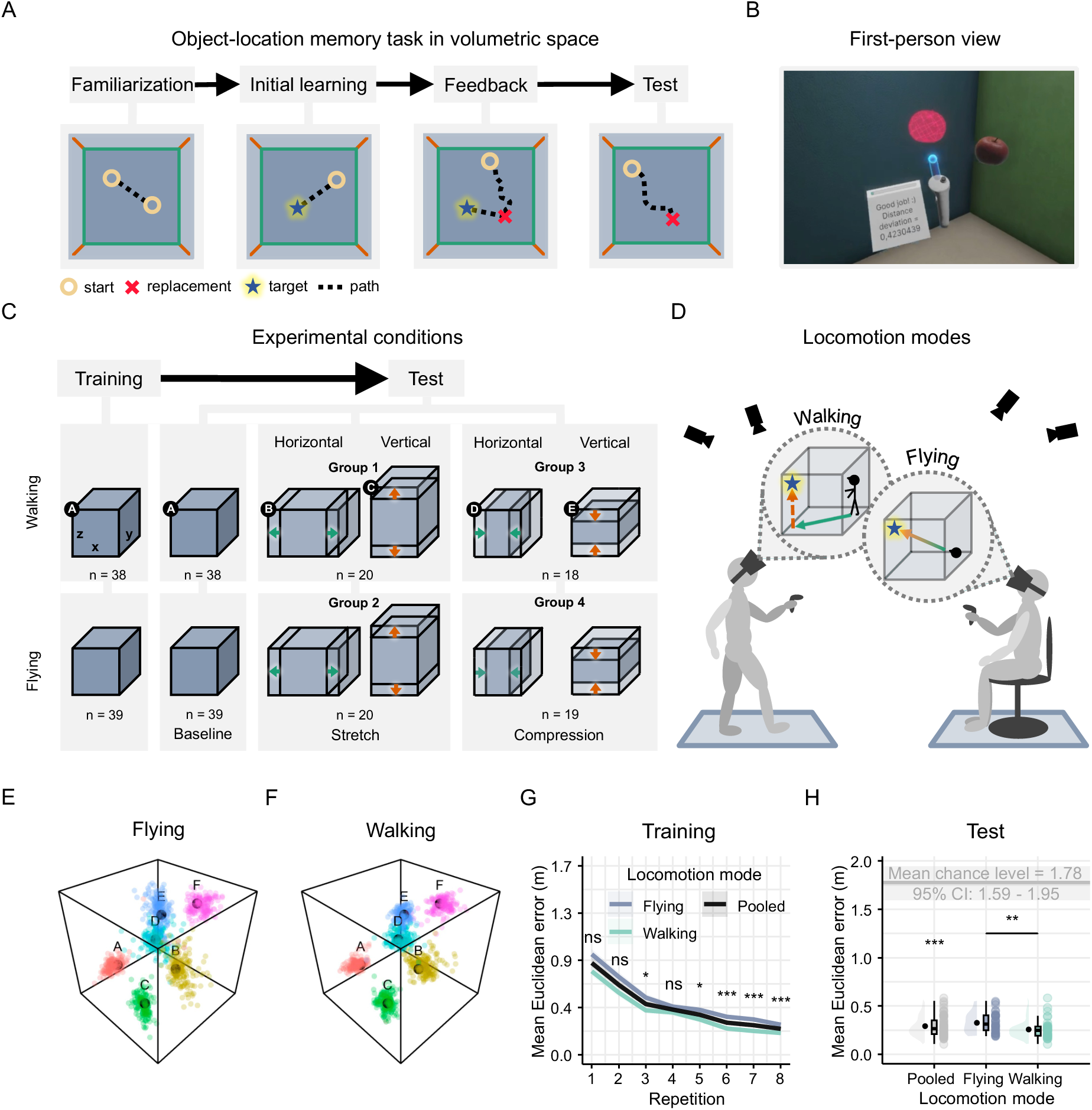
Experimental task and design. **A:** Object-location memory task in volumetric space. *Familiarization*: Participants practiced navigating (dotted line) within the 3D virtual environment by collecting freely-floating balls (circles) in succession. *Initial learning*: Participants memorized virtual everyday-life objects at six target locations, fixed in volumetric space (star), always starting from random locations (indicated by a ball). *Feedback*: Participants replaced each object at its corresponding target location from memory (cross), starting from a ball’s random location. After each response, they received immediate feedback about their accuracy and re-encoded the object’s correct location before the beginning of the next trial. *Test*: Participants continued replacing objects at remembered target locations but without receiving feedback. **B:** First-person view during feedback. Feedback was provided based on the displacement of the object replacement from the target location in meters. The true target location was displayed alongside the corresponding replacement location (red hologram). **C:** During training, object-locations were learned in the baseline environment (cube). During the test, spatial memory was assessed in both the baseline and in the deformed environments. Conditions were manipulated in a 2 × 2 × 2 mixed design with the within-subject factor *environmental dimension* (horizontal vs vertical plane) and the between-subject factors *type of deformation* (stretched vs compressed) and *mode of locomotion* (flying vs walking). The baseline environment (3 × 3 × 3 m) was deformed by a factor of 1.33 (33%) for horizontal or vertical dimensions (see Supplemental Tab. S2). Note that all participants were naive about the deformations. **D:** We divided participants into two groups with different modes of locomotion. In the walking group, participants freely moved on the horizontal surface of the enclosure and they reached vertical positions by extending their arm or a virtual wand. In the flying group, participants virtually navigated in all three dimensions while physically sitting on a rotatable chair. Translation movement (forward/backward) was achieved via the controller, and flying direction (pitch × yaw) was set by their head rotation. Motion-capture cameras recorded the movements of a set of rigid bodies attached to participants’ body parts (see Supplemental Fig. S1B). **E, F:** Replacement locations (color-coded) and the true target locations (labeled black dots), separately for the walking (E) and flying (F) group in the baseline condition. Each dot represents a single trial object replacement of a given participant during the test phase. **G:** Baseline training performance (Euclidean error in 3D space) as a function of repetition, averaged across target locations, separately for each locomotion mode and pooled (color-coded lines depict means ± SEM). Mean spatial memory error (in meters) decreased over repetitions. Flying participants showed higher errors than walking participants in the last few trials. **H:** Baseline test performance (Euclidean error in 3D space) for each locomotion mode and pooled. The mean spatial memory error was significantly lower than the mean chance level (gray horizontal line) with flying participants showing higher errors than walking participants. Violin plots depict the density distribution, boxplots the median and quartiles, black dots with error bars the means ± SEM, and colored dots individual data points per condition. * *P* < .05, ** *P* < .01, *** *P* < .001.

### Successful encoding of 3D coordinates in volumetric space

Previous research in humans investigated spatial memory in 3D space exclusively by probing target locations projected onto surfaces or within discrete spaces. But can human participants sufficiently encode object-locations in fully volumetric space? To test this, we assessed participants’ spatial memory accuracy during training and test (environment A in Fig. 1C; Fig. 1E-F). For each trial, we measured the distance between the object-replacement and the correct target location in 3D Euclidean space indicating a mnemonic error in representing object-locations. During training, spatial memory errors overall decreased as a function of replacement repetition (1-way repeated-measures ANOVA: *F*2.93,222.3 = 122.76, *P* < .001, 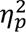 = .618; Fig. 1G) with an average improvement of .6 m from the first to the last repetition (Post-hoc paired *t*-test: *t*76 = 15.3, *P*adj < .001, *d* = 1.74; Bonferroni-corrected for 28 comparisons). During test (when no feedback was provided), spatial memory accuracy remained high (M = .30 m, SD = .10 m), significantly falling below the mean chance level based on simulated random errors (M = 1.77 m, SD = .24 m, bootstrapped 95% CI = 1.60-1.94 m; 1-sample *t*-test: *t*76 = -127, *P* < .001, *d* = -14.4; Fig. 1H), demonstrating that participants successfully encoded freely floating objects in volumetric space. However, when comparing spatial memory performance across different modes of locomotion, we found that flying participants exhibited overall higher errors than walking participants, both at training (1-way ANOVA: *F*1,614 = 20.811, *P* < .001, 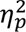 = .033; Fig. 1G) and at test (2-sample *t*-test: *t*73.4 = 3.07, *P* = .003, *d* = .7; Fig. 1H).

### Anisotropic representation of volumetric space is locomotion dependent

To evaluate whether humans memorize volumetric space equally across spatial dimensions and how this might be influenced by the mode of locomotion, we analyzed dimension differences in participants’ response dispersion. To this end, we pooled responses across target locations for each group (Fig. 2C/E), for target-specific responses see Fig. 2A-B; Supplementary Fig. S2) and computed the median absolute deviation (MAD) of responses (Fig. 2D/F). A significant interaction between spatial dimension and locomotion mode was found in the baseline environment (2-way mixed ANOVA: *F*1,75 = 40.73, *P* < .001, 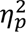 = .352), with a higher response dispersion along the vertical compared to the horizontal dimension in the flying group (paired *t*-test: *t*36 = 4.34, *P* < .001, *d* = .713, Bonferroni-corrected for 2 comparisons; Fig. 2D). A contrary effect was observed in the walking group (*t*39 = -4.99, *P* < .001, *d* = -.789; Fig. 2F), suggesting a lower vertical precision for flying participants and a higher vertical precision for walking participants. Notably, we found a significant difference between the flying and walking groups only in the vertical dimension (*t*55.8 = 5.88, *P* < .001, *d* = 1.36), but not in the horizontal dimensions (*t*75 = .698, *P* = .974, *d* = .159), indicating that only vertical information is represented differently depending on the mode of locomotion.

**Figure 2.**
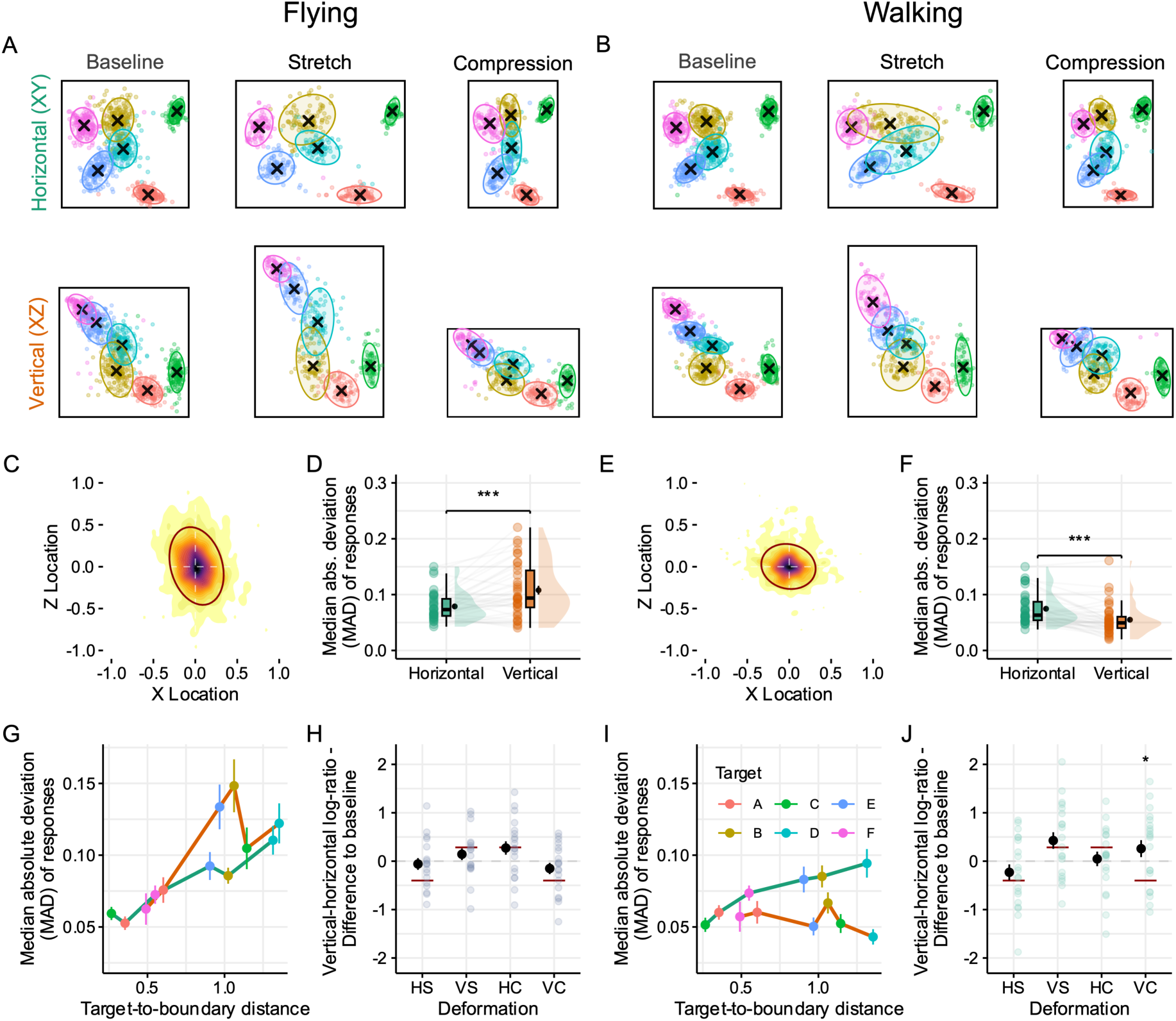
Locomotion-dependent anisotropy in the representation of volumetric space. **A, C-D, G-H:** Flying group. **B, E-F, I-J:** Walking group. **A, B:** Trial-wise object replacements projected onto 2D planes, deformation type (columns) and deformation axes (rows). Ellipses refer to areas including 95% of responses for different target locations (color-coded) with black crosses demarking their median. **C, E:** Distribution of median-centered replacement responses in the baseline testing environment, pooled over target locations, reveals larger dispersion along the z-axis in the flying condition (C) and larger dispersion along the x-axis in the walking condition (E). Darker colors reflect higher frequency of values. **D, F:** Median absolute deviation (MAD in meters) in the baseline for the horizontal (x-,y-axis) and vertical (z-axis) for the flying (D) and walking (F) groups. Violin plots depict the density distribution, boxplots the median and quartiles, mean ± SEM as black dots with error bars, as well as individual data points (dots) per condition. *** *P* < .001. **G, I:** MAD for different target locations as a function of target-to-boundary distance, separately computed for the horizontal dimension (green line) and vertical dimension (orange line). While the flying group (G) shows a positive distance-MAD relationship in both dimensions, the walking group (I) only shows this relationship in the horizontal dimension. **H, J:** Vertical anisotropy indices defined as vertical to horizontal MAD log-ratio for different deformation types (HS/VS: Horizontally/vertically stretched, HC/VC: Horizontally/vertically compressed) relative to baseline. Participants’ ratios were tested against the expected elongation based on the scaling factor of the deformation environments (red bars; stretch: 1.33, compression: .67). VC condition in walking group differed significantly from the expected elongation. * *P* < .05 (Bonferroni-corrected for 8 comparisons).

Next, we examined whether differences in navigational behavior between locomotion modes underlie the observed differences in vertical processing. We found that participants of the walking group visited fewer locations in the volumetric space compared to the flying group (2-sample *t*-test: *t*74.5 = 3.38, *P* = .001, *d* = .771; Supplementary Fig. S4C) and their movement paths between the start and replacement location were shorter (*t*71.8 = -2.24, *P* = .028, *d* = - .508), indicating higher confidence in spatial memory. Their location-specific movement paths were also much more similar across trials (*t*61.7 = 10.1, *P* < .001, *d* = 2.32), suggesting stronger reliance on route-based strategies. Moreover, their head orientation at replacement during test differed less from the orientation at collection during training than in the flying condition (*t*52.4 = 3.57, *P* < .001, *d* = .823), indicating a stronger tendency for perceptual matching. Only heading difference (Spearman’s *Rho* = .52, *S* = 36594, *P* < .001; Supplementary Fig. S4D) and path dissimilarity (Spearman’s *Rho* = .66, *S* = 25672, *P* < .001; all other *P* > .126) correlated with vertical dispersion. Together, these results highlight the role of the body serving as an egocentric reference frame during naturalistic, bipedal walking that can shape the vertical-horizontal symmetry of human spatial memory.

### Distinct use of vertical boundaries and body cues for height estimation during naturalistic walking

In 2D rectangular environments, locations are encoded relative to the enclosure’s boundaries, resulting in better memory for locations close to boundaries than far from them (Hartley et al., 2004; Lee et al., 2018). To evaluate whether the same relation holds for both horizontal and vertical boundary in each locomotion mode, we computed the correlation between the response dispersion and the distance to the nearest boundary for each dimension (Fig. 2G/I). As expected, both locomotion groups exhibited a comparable positive relationship between the response dispersion and the target-to-boundary distance in the horizontal dimension (walking: average Spearman’s *Rho* = .33, *t*39 = 4.84, *P* < .001; flying: Spearman’s *Rho* = .481, *t*36 = 4.84, *P* < .001). However, the two groups differed significantly with respect to the vertical dimension. Flying participants showed a positive (Spearman’s *R*ho = .327, *t*36 = 5.27, *P* < .001; Fig. 2G) and walking participants an absent or even reversed relationship (Spearman’s *Rho* = -.0429, *t*39 = -.551, *P* = .584; Fig. 2I). In other words, the location that has the largest target-to-boundary distance, approximately equidistant between the floor and ceiling (target location D in Fig. 2I) exhibited the highest vertical dispersion for flying participants and the lowest for walking participants (flying: M = .122, SD = .085; walking: M = .0431, SD = .0339; 2-sample *t*-test on walking vs. flying: *t*46.5 = 5.28, *P* < .001).

One possible explanation for why walking participants might process vertical spatial information differently from flying participants is that, in addition to relying on environmental boundaries, they could encode locations relative to their upright body. This posture is reliably anchored to the plane of locomotion and aligned with gravity. Thus, locations near the vertical center are not only farthest from vertical boundaries but also closest to the participant’s head height (participant height: M = 1.74 m, SD = .07 m; height of object-location D: 1.65 m). Indeed, we found increased spatial memory dispersion as the distance between the height of a target location and the height of the walking participant’s head increased (1-sample *t*-test on individual correlations between MAD and target-to-head distance: Average Spearman’s *Rho* = .18, *t*39 = 2.26, *P* = .029). This suggests that both vertical boundaries and the participants’ own bodies, particularly the position of their heads, serve as useful reference points for estimating height during naturalistic walking.

To investigate the role of boundaries in spatial memory for volumetric space in more detail, we analyzed how objects are replaced in test environments that have undergone geometric deformations. For instance, if the cubic baseline environment was stretched along one dimension, it is reasonable to expect a greater dispersion of responses along the same dimension, but not along the orthogonal (unchanged) one, potentially amplifying or reducing baseline anisotropies. The mode of locomotion could further influence changes to isotropy due to differences in environmental boundary processing between flying and walking. We computed anisotropy indices as the log-ratio between the vertical and horizontal MAD for all deformed environments and calculated difference scores with respect to the baseline environment (deformation ratio - baseline ratio; Fig. 2H/J). We then tested whether the anisotropy indices changed in the expected direction and according to the scaling of the deformed environment (1.33 for stretch, .67 for compression). Participants’ response dispersion, indeed, changed in a dimension-specific manner approximating the scaling of the deformed environments. There were no significant differences to the expected values, except for the vertical compression condition when participants were walking (1-sample *t*-test against scaling-adjusted difference to baseline: *t*19 = 3.76, *P* = .011, *d* = .840, all other tests: *P* > .094; Bonferroni-corrected for 8 comparisons). Here, the dispersion increased rather than decreased as expected from the compression, indicating that vertical compression heightened participants’ uncertainty, likely because a ground-based encoding strategy became less effective. For instance, if a target location’s height was initially far above the head and encoded relative to the ground in the baseline environment, vertical compression of the environment would force participants to adopt a novel reference strategy.

### Flying participants employ a boundary-proximity strategy across horizontal and vertical deformations

Manipulating the size and shape of a familiar task environment has been previously conducted in 2D environments to investigate the causal impact of environmental geometry on cognitive maps across species (O’Keefe & Burgess, 1996; Barry et al., 2007; Stensola et al., 2012; Keinath et al., 2018, 2021; Hartley et al., 2004; Zisch et al., 2024). To evaluate how object-locations are encoded in volumetric space, we extended a set of geometric models (Hartley et al., 2004) from 2D to 3D space and tested their predictions on object replacements in the deformed environments (Fig. 3A). Each model assumes a different encoding mechanism of the object-location in the familiar baseline environment and predicts object replacements in deformed test environments that preserve their respective encoding mechanism. While the *fixed distance* and *fixed ratio* models preserve either the distances to the nearest walls or the aspect ratios to opposing walls, respectively, the *boundary-proximity* model encodes the proximity to all surrounding walls in a non-linear fashion. Specifically, the model assumes that locations close to boundaries are encoded more by their fixed distance and locations in the center more by their fixed ratio (see Methods section under “Modeling”). Consequently, the boundary-proximity model mimics place predictions similar to those derived from the population activity of boundary vector cells (BVCs) each tuned to a preferred allocentric direction and distance to walls (Hartley et al., 2000; Barry et al., 2006). We first fitted the model parameters to the object replacements of the baseline environment to account for baseline differences in precision to the three spatial dimensions, and then computed the model-likelihood of replacements in the deformed environments. Across all participants, the object replacements were best explained by the boundary-proximity model (1-way repeated-measures ANOVA: *F*1.18,89.96 = 108.963, *P* < .001, 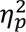 = .589; post-hoc paired *t*-test, Bonferroni corrected for 3 comparisons: proximity vs. ratio: *t*76 = 7.71, *P* < .001, *d* = .879; proximity vs. distance: *t*76 = 11.8, *P* < .001, *d* = 1.35; ratio vs. distance: *t*76 = 9.15, *P* < .001, *d* = 1.04). We further found a significant 3-way interaction between the model, mode of locomotion and the dimension of deformation (3-way mixed ANOVA: *F*1.16,87.33 = 14.459, *P* < .001, 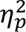 = .162). Post-hoc paired *t*-tests confirmed the superiority of the boundary-proximity model (proximity model against other models: all *P* < .012, Bonferroni-corrected for 12 comparisons; see Fig. 3B), suggesting the involvement of BVC-like computations in spatial memory during 3D deformations of the environment. Similar to previous finding in 2D environments (Hartley et al., 2004), the relative fit of the fixed ratio model vs. the fixed distance model was higher for the targets that were farther away from the wall in all conditions, except for the walking group in the vertically stretched environment (Fig. 3D & E). In general, responses to vertical deformations revealed a lower fit to boundary-proximity model predictions in the walking, but not in the flying group (2-way mixed ANOVA: *F*1,75 = 46.088, *P* < .001, 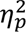 = .381; horizontal vs vertical for walking: *t*39 = 7.15, *P* < .001, *d* = 1.13; flying: *t*36 = -.337, *P* = 1, *d* = -.055; paired *t*-test, Bonferroni-corrected for 2 comparisons).

**Figure 3.**
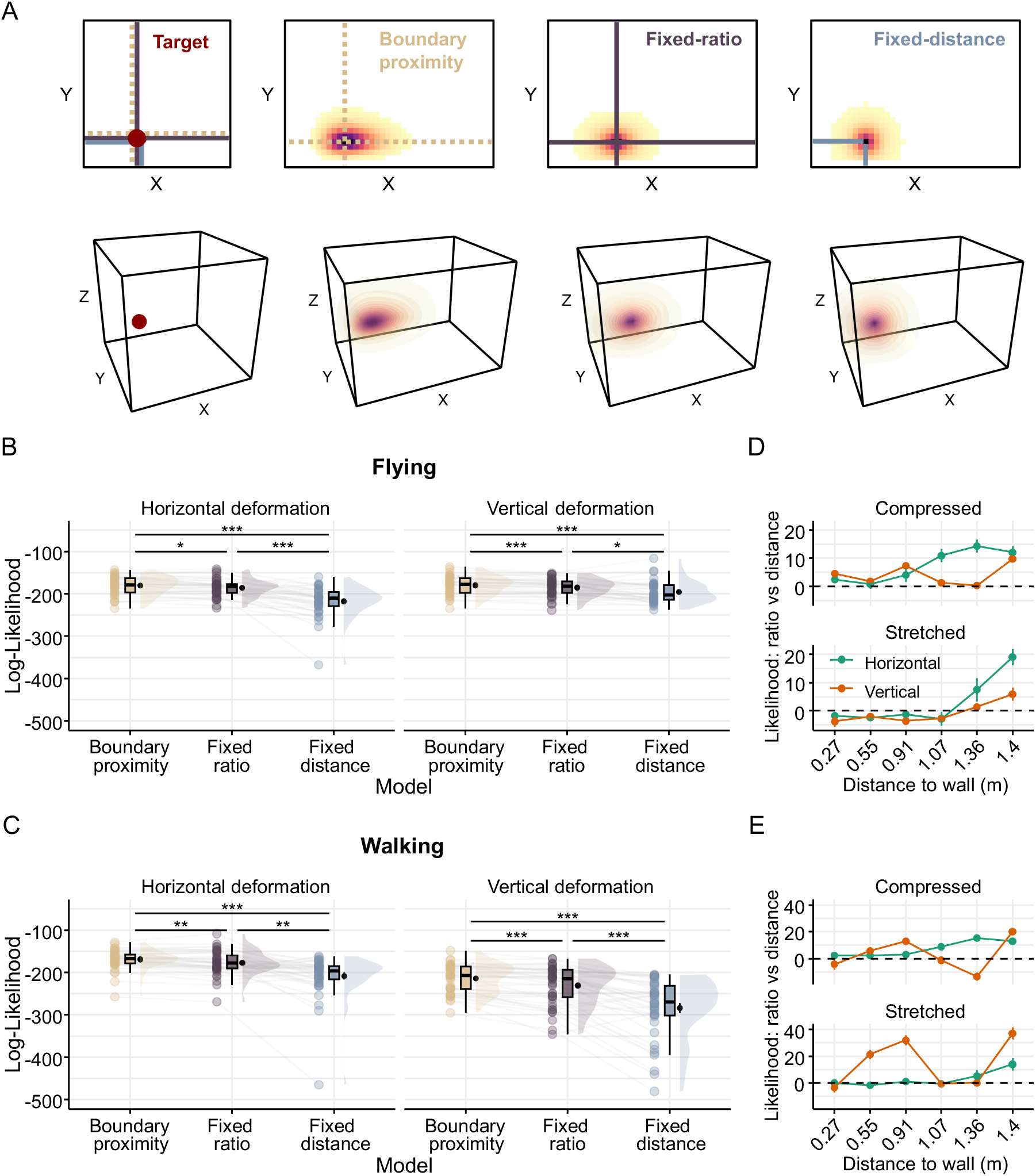
Fitting geometric models to participants 3D object replacements. **A:** Three geometric models applied to the object replacement data: Fixed distance model (preserves the fixed distance of a memorized location to the nearest two walls), fixed ratio model (preserves aspect ratios between opposing walls), boundary-proximity model (preserves the proximity - inverse of non-linear distance - to each of the walls). Heat maps represent the probability of an object replacement for a given target location from each model prediction (left, red dot). Top: 2D projection of the probability matrix for the x- (stretched) and y-axis (non-stretched). Bottom: 3D-extended prediction volumes. **B, C:** The boundary-proximity model shows the best fit to the replacement data across conditions. Displayed are log-likelihood model fits for each model, separately for horizontal and vertical deformations and for the flying (B) and walking (C) condition. Violin plots depict the density distribution, boxplots the median and quartiles, black dots with error bars the means ± SEM, and colored dots individual data points per condition. * *P* < .05, ** *P* < .01, *** *P* < .001 (Bonferroni-corrected for multiple comparisons). **D, E:** Replicating previous findings in 2D space (Hartley et al., 2004), we observe a relationship between 3D distance of a target location to the walls (x-axis) and the relative fit (log-likelihood, y-axis) to the fixed ratio vs fixed distance model for different deformation types. This is especially the case for horizontal compared to vertical deformations (color-coded). The vertical stretch deformation in the walking group deviates from this pattern.

### Walking participants rely on a ground-based strategy in vertical deformations

Consistent with our findings in the baseline environment, the explanatory power of the geometric models decreased for vertical deformation in the walking participants. Contrary to the flying group, walking participants maintained an upright posture aligned with the gravitational axis, allowing them to estimate the height of objects from the ground by using their body as a reference. If this is true, vertical coordinates of objects should be encoded with greater reference to the ground than to the ceiling (cf., Fig. 2A). Accordingly, we tested a modified version of the boundary-proximity model that preserves the proximity to all walls except the ceiling (*ground-proximity* model), thereby amplifying the weight on the influence of the ground (Fig. 4A). Our analysis revealed a significant interaction between the model type and locomotion mode for the vertical deformations (2-way mixed ANOVA: *F*1,75 = 28.876, *P* < .001, 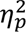 = .278; Fig. 4B). The ground-proximity model provided a better fit for data from walking participants compared to the standard boundary-proximity model (paired *t*-test: *t*39 = 2.86, *P* = .014, *d* = .452; Bonferroni-corrected for 2 comparisons). In contrast, data from flying participants was better explained by the standard boundary-proximity model (*t*36 = -11.0, *P* < .001, *d* = -1.81).

**Figure 4.**
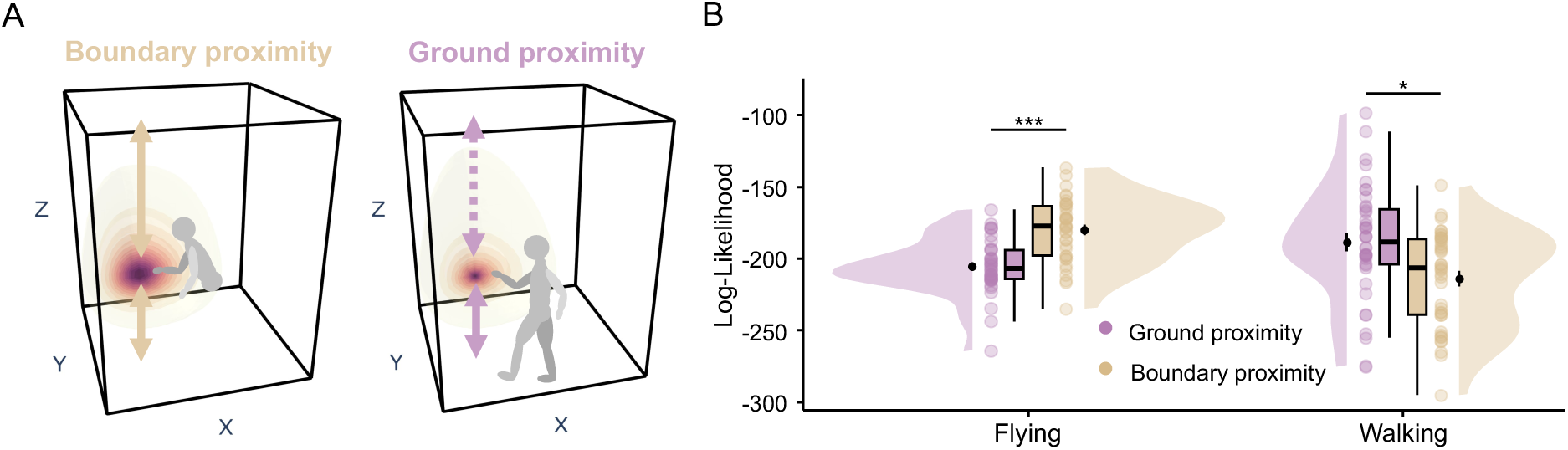
Locomotion-dependent use of vertical boundaries. **A:** Schematic of a boundary proximity vs ground proximity prediction. Unlike flying participants (*left*), walking participants (*right*) are anchored to the ground and maintain an upright posture. Thus, they might put more emphasis on the ground than the ceiling when encoding vertical coordinates. We modified the boundary-proximity model (yellow arrows and prediction field, left) by removing the reference to the ceiling and termed it ground proximity model. **B:** Model comparison between the ground-proximity model and the standard boundary-proximity model in each locomotion group. The ground-proximity model showed a superior fit (high log-likelihood) for the walking group, but not for the flying group. Boxplots depict the median and quartiles, black dots with error bars the means ± SEM, and colored dots individual data points per condition. * *P* < .05, *** *P* < .001 (Bonferroni-corrected for two comparisons).

## Discussion

The present study employed immersive Virtual Reality (VR) to investigate the mechanisms underlying the representation of volumetric space, focusing on how different modes of locomotion (flying vs walking) and environmental geometry (cubic vs. horizontally and vertically deformed environments) affect spatial memory.

The significance of environmental boundaries for spatial memory has been well-established on a two-dimensional horizontal plane. Previous research shows that humans recall locations near boundaries with greater precision than those farther away (Hartley et al. 2004, Lee et al. 2018), which has been linked to the firing properties of boundary vector cells (BVCs; Barry et al., 2006; Hartley et al., 2000) initially found in the rat subiculum (Lever et al., 2009). The firing of BVCs is tuned to preferred distances and directions to environmental boundaries with receptive fields that broaden as the distance from these boundaries increases, indicating diminished encoding precision. Our study suggests a similar mechanism for the encoding of volumetric space when participants were flying virtually. In the baseline environment, we observed that spatial memory precision peaked when target locations were close to any boundary — be it the ceiling, floor, or side walls. Moreover, when the test environment was vertically or horizontally deformed, flying participants adjusted their spatial memory in ways predicted by a 3D-extended boundary-proximity model which was inspired by putative 3D BVCs tuned to both vertical and horizontal boundaries. These findings imply that humans use boundaries to represent 2D and 3D space in a qualitatively similar manner when movement constraints are minimal — much like birds or fish exploring their natural habitats.

However, our data suggest that spatial encoding is not isotropic in volumetric space. Flying participants showed a lower encoding precision for the vertical compared to the horizontal dimension, contrasting with findings in flying bats, which demonstrate isotropic place fields (Yartsev & Ulanovsky, 2013). This anisotropy could stem from the spatial ecology during ontogeny and evolution of surface-dwelling animals, including biases in sensory processing, energy costs of moving against gravity, and the availability of spatial cues (Davis, Holbrook & de Perera, 2018). Specifically, visual perception may be less precise when the head is tilted up or down compared to when it is oriented straight ahead (Dhaliwal et al. 2002, Skerswetat et al. 2020). In addition, it is likely that the method of flying in our study (i.e. moving virtually via joystick while seated) introduced a mismatch between visual and proprioceptive feedback. Such a sensory conflict could result in heterogeneous or inconsistent responses in spatially tuned cells (Chen et al., 2013) and may disrupt spatial processing, especially in the vertical axis, where physical gravity imposes a constant influence. This raises the intriguing question of whether anisotropy would persist in environments with near-zero gravity, such as on space stations (Knierim et al., 2000). In these settings, anisotropy might be undetectable due to the equivalence of all spatial axes, or it might emerge individually, depending on the subjective anchoring of “up” and “down” as reported by astronauts (Oman, 2007).

Anisotropy in volumetric spatial encoding may also be reflected at the neural level. One possibility is that BVCs can have preferred directions to any boundary but have larger tuning widths for vertical boundaries compared to horizontal ones. Alternatively, there may be differing numbers of BVCs tuned specifically to vertical or horizontal boundaries, akin to the distribution of head direction cells in bats, where cells tuned to horizontal directions are more abundant than those tuned to vertical ones (Finkelstein et al., 2015). Supporting this, human fMRI research has identified distinct neural structures involved in encoding vertical and horizontal directions during virtual flying (Kim & Maguire, 2019). More recently, BVCs tuned to vertical or horizontal walls were detected in goldfish swimming in a quasi-planar water tank (Cohen et al., 2023). Electrophysiological recordings in flying or swimming species within bounded volumetric environments could further elucidate whether BVCs are tuned to all three axes with varying tuning widths or are selectively tuned to specific planes.

In contrast to the flying condition, boundary-dependent 3D memory exhibited significant differences when participants walked, constrained to the ground plane. Here, precision for horizontal locations remained highest near boundaries, but vertical precision peaked near the center of the environment, aligning with participant’s head positions. The head, containing critical sensory organs such as the eyes for visual and the inner ear for vestibular input, provides essential cues for vertical estimation. During naturalistic walking, humans’ upright posture and gravity-aligned body axis serve as an intrinsic “ruler” for estimating vertical locations, supplementing environmental boundaries. This reliable vertical cue could compensate for the lower vertical precision in the center of an environment. In contrast to quadrupedal animals, bipedal humans can access a broader range of vertical space. Various body parts, including the eyes, head, trunk and limbs, can serve as useful reference frames for volumetric space, putatively integrated into body-centered coordinates within the posterior parietal cortex (Andersen, Essick & Siegel, 1985; Buneo et al., 2002; Cohen & Andersen, 2002; Whitlock, 2017). The abundance of body-based information under gravity may have encouraged participants to adopt more egocentric strategies, such as perceptual matching between encoding and retrieval phases or route-based navigation, as observed in the walking group. Enhancing the degree of embodiment by incorporating a visible virtual avatar could further amplify precision for vertical locations by providing additional egocentric reference cues. Future research could investigate naturalistic walking scenarios with varying levels of avatar representation — ranging from showing hands and arms to the entire torso — to better understand the role of body-based cues in spatial memory for vertical locations. It is important to note, however, that increased vertical precision might diminish or even reverse for targets located outside of peripersonal space, similar to the patterns observed in the flying group.

The strong influence of gravity and body references during walking necessitated modification of boundary-based computational model when predicting spatial memory in deformed environments. Unlike the 3D boundary-proximity model, which provided the best fit to the flying group’s data, the ground-proximity model which referenced only the floor but not the ceiling — best explained the walking participants’ memory. For crawling or walking animals, posture relative to the ground is a salient cue for spatial memory. Studies have shown that rodents, humans, and birds can distinguish locations on flat versus tilted surfaces, aiding their navigation and spatial memory (Groberty & Shenk, 1992; Steck et al., 2003; Nardi & Bingman, 2009).

Our findings emphasize the complementary roles of environmental boundaries and the gravity-aligned body axis in representing 3D space. Cognitive maps exhibit flexibility, adapting to multiple factors, including the mode of locomotion, attention, and degree of embodiment. For instance, a volumetric map may be constructed when flying, whereas a quasi-planar map may be utilized when walking (Finkelstein et al., 2016; Kim & Doeller, 2022, 2024). Attentional shifts between local surface cues (e.g., floor texture) and global cues (e.g., environmental boundaries extending into 3D space) likely influence the representational format.

In conclusion, our study demonstrates that cognitive maps of volumetric spaces share a common boundary dependency with those of two-dimensional environments. However, differences in locomotion modes introduce significant variations in the use of boundary and body-based references. As terrestrial, bipedal navigators, humans encode the axis of gravity differently from the plane of locomotion, using their bodies as “rulers” to measure distance from the ground. The finding that gravity-defying conditions within the same species result in distinct vertical representations and reference frames highlights the flexibility of the cognitive mapping system, which is adapted for navigating a complex, multidimensional world.

## Methods

### Participants

A total of 85 young healthy adults were recruited from the internal participant database of the Max Planck Institute for Human Cognitive and Brain Sciences in Leipzig, Germany. The sample size was determined by a power calculation using G*Power version 3.1.9.6 (Faul, Erdfelder, Buchner & Lang, 2009). We assumed a small-to-medium effect size of *f^2^* = .23 (directly derived from partial η^2^ = .05) for the interaction effect between 4 groups (flying vs walking locomotion mode, and stretch vs compression deformation) and 2 repeated measures (horizontal and vertical deformation) tested by a repeated measures ANOVA assuming non-sphericity and a medium correlation among within-subject factors of r = .4. This resulted in a sample size of 17 participants per group (*N* = 68) to achieve a statistical power of 80%, with an error probability of α = .05 (two-tailed test). Data from 14 additional participants were acquired to account for potential dropouts. 3 participants did not complete the experiment due to motion sickness or fatigue. Data from 5 participants were excluded due to poor spatial memory performance at baseline, classified as having mean spatial memory errors greater than 1.5 times the interquartile range above the upper quartile of mean errors observed in the sample. Thus, data from 77 participants (40 female, 36 male, 1 non-binary; Age: 26.182 ± 4.468 years, 19-35 years) entered the analysis. All participants were right-handed, had normal (or corrected-to-normal) vision, and no current or past neurological, psychiatric, or motor disorder. They were reimbursed for their participation, following institutional procedures. To minimize the variability in spatial abilities and movement dynamics due to differences in body height, we included only participants with a height between 165 cm and 185 cm. All participants gave informed consent in accordance with the Ethics Advisory Board of the Medical Faculty of the University of Leipzig. Participants were pseudo-randomly assigned to 4 experimental groups of approximately equal size, with no significant differences in demographic variables and body measures between groups (all *P*’s ≥ .08). A complete record of group-specific sample characteristics is provided in Supplementary Table 1.

### Experimental setup

The experiment took place in a Virtual Reality (VR) lab at the Max Planck Institute for Human Cognitive and Brain Sciences in Leipzig, Germany (Supplementary Fig. S1A). Participants wore a Pico Neo 3 Pro Eye HMD (3664 × 1920 pixels, horizontal field of view: 98 degrees, vertical field of view: 90 degrees, refresh rate: 90 Hz, weight: 620 g; Pico Interactive, San Francisco, CA, USA https://www.picoxr.com) displaying custom-made virtual environments created in Unity3D (version 2021.3.3f1; Unity Technologies, San Francisco, CA, USA) and rendered on a stationary computer (AMD Ryzen 5 3600 6-core 3.6-4.2 GHz processor, NVIDIA GeForce RTX 3090 graphics card, Intel I350 network card, running Microsoft Windows 10). Within the virtual environment, the participants’ viewpoint was continuously updated based on their real-time head location and movement in virtual space, allowing for naturalistic behavior. Full-body movements were recorded at a sampling rate of 200 Hz (downsampled to 40 Hz) using a Vicon MoCap system (Vicon, Oxford, UK; https://www.vicon.com/) with 9 Vero v2.2 infrared cameras at a resolution of 2.2 MP covering a tracking volume of approximately 4 × 5 × 2.5 m. During the experiment, participants wore passive retro-reflective markers attached to 13-14 rigid bodies, the HMD, and a handheld motion controller (Supplementary Fig. S1B). 3D motion data was live-streamed into the SteamVR-based application, which maintained a wireless connection to the HMD via the PCVR streaming software Pico Link and a Wi-Fi 6 router (ASUS RT-AX82U). The VR application was remotely controlled and monitored by the experimenter.

A follow-up debriefing questionnaire was created using SoSciSurvey version 3.5.01 (Leiner, 2024; www.soscisurvey.de) and presented on a standard computer screen (2560 × 1440 pixels).

### Study design and virtual environments

We implemented a 2 × 2 × 2 mixed factorial design manipulating the within-subject factor spatial dimension (horizontal vs. vertical) and the between-subject factors deformation type (stretch vs. compression) and locomotion mode (walking vs. flying). An overview of the experimental conditions is provided in Figure 1C.

During training, participants learned the 3D location of multiple virtual objects within a symmetrically shaped cubic baseline environment. During the test, their spatial memory was probed both in the familiar environment and in geometrically deformed environments (see “Object-location memory task in 3D space” under “Experimental protocol”). While for one group of participants the test environment was stretched (groups 1 and 2), for another group it was compressed (groups 3 and 4). For all participants, both the vertical and horizontal dimensions of the environment were deformed separately. The task was performed in two modes of locomotion (Fig. 1D). One group of participants physically walked on the ground, reflecting natural human locomotion (groups 1 and 3). Another group of participants was able to fly virtually through space, mimicking the natural locomotion mode of winged animals (group 2 and 4). These participants sat in a motion-captured rotatable chair and controlled their 3D locomotion direction by rotating their head (yaw & pitch) and forward/backward translation using the controller’s joystick. The maximum speed of movement was constant across all environments at 1 m/s, reached after .5 s after the initial joystick push/pull, with a smooth acceleration at a rate of 1.5 m/s^2^. All participants could reach positions within peripersonal space by extending the arm upwards or bending downwards. Participants in the walking group were able to extend the virtual arm controller like a wand if a target could not be reached by physically extending the arm upwards (controller position ≥ 15 cm above the head and forward tilt angle of ≤ 55 degrees).

Spatial memory was assessed in 5 different virtual environments, one baseline cube and four deformed cuboids (horizontally stretched, horizontally compressed, vertically stretched, vertically compressed). The baseline environment consisted of a cubic enclosure (3 × 3 × 3 m) made of six equally sized walls. These walls were given unique colors and tile textures to facilitate orientation and provide optical flow during locomotion (Supplementary Fig. S1G). The assignment of colors and textures to walls was randomized across participants but fixed throughout the experiment. The deformed environments were exact duplicates of the baseline environment, except that they were uniformly stretched or compressed by a fixed rescaling factor of .33 (approximately 1 m) along either the horizontal (x) or vertical (z) axis, relative to the baseline environment. All virtual environments were symmetrically illuminated, so that light could not be used as a directional cue. No local landmarks were provided at any time to enforce spatial learning based on the geometry of the environment. Detailed characteristics of the virtual environments are shown in Supplementary Table. 2.

### Experimental protocol

Participants completed the following steps in consecutive order: preparation, main task, and debriefing. The entire experimental session took 60-210 min (M = 148 min; SD = 30 min) with 42-162 min (M = 84 min, SD = 24 min) in VR.

#### Preparation

Upon arrival, participants read the written task instructions and were informed about the general objectives, procedure and potential risks (e.g., cybersickness) participating in the study. After giving consent, the experimenter recorded several body measurements (body height, arm span, inseam length, hip width and arm length) and attached the rigid bodies to the participants’ body, with the configuration varying slightly between conditions (Supplementary Fig. S1B). Participants did not wear shoes throughout the experimental session. We then fitted a virtual avatar to the participant’s T-pose to scale a simple digital 3D model of the participant’s body based on individual measurements. Importantly, the avatar was not visible to the participant during the entire experiment. All participants started the experiment from the center of the lab’s horizontal plane, either standing or sitting in a chair. Throughout the session, condition-specific task instructions were delivered via a virtual display attached to the animated controller (Fig. 1B).

### Object-location memory task in volumetric space

Participants performed an object-location memory task, which we extended from 2D to 3D space. The general task structure followed Doeller et al. (2008) and can be divided into four successive phases: familiarization, initial learning, feedback, and test (Fig. 1A). Exemplary task trials for both locomotion modes are shown in Supplementary Movie 1-4 (https://osf.io/r7ubv/).

#### Familiarization

(Supplementary Movie 1). Participants were spawned into the virtual environment and practiced task-relevant interactions and movements during the familiarization phase. In each trial, they were instructed to move freely to the floating green ball and collect it by pointing the tip of the animated controller into the ball until it lights up and then pressing a button. After collection, the ball reappeared at a different location within the virtual environment. Participants completed 6 familiarization trials (duration: M = 1.8 min, SD = 1.2 min).

#### Initial learning

(Supplementary Movie 2). In each trial of the subsequent initial learning phase, participants moved to a start location, indicated by a green ball. Once collected, the ball disappeared and a target object was displayed at its predefined location. Participants were asked to move to the object, memorize its location and collect it. There were 6 trials, one for each target location (duration: M = 3.0 min, SD = 2.4 min).

#### Feedback

(Supplementary Movie 3). Next, learning was continued through feedback-based training. On each trial of the feedback phase, participants again collected a green ball at a 3D start location. A 3D model of the object then appeared attached to the controller, prompting them to place the object at the remembered location. At the same time, the name of the object was printed on the controller’s display. Once the response was confirmed by pressing a button (“response”), its location was marked in the form of a hologram as the object moved to its correct location. In addition, accuracy-related feedback was printed on the controller’s display via one of five smileys and a numerical error value. The feedback was based on the 3D Euclidean distance between the response and the correct location, using predefined thresholds (0-.17 m = very good, .17-.27 m = good, .27-.47 m = intermediate, .47-1.06 m = bad, > 1.06 m = very bad). These thresholds were derived via 3*D Euclidean error* 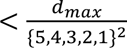, where *D_max_* represents the maximum possible distance in a 3 × 3 m plane. Finally, participants recollected the object before the next trial began. In total, there were 48 trials with 8 repetitions per target location (duration: M = 27.6 min, SD = 10.8 min).

*Test* (Supplementary Movie 4). Ultimately, spatial memory was probed during the test phase, which served as a readout of spatial memory, i.e., the result of training. Trials in the test phase followed the same timeline as the feedback trials, except for feedback and re-encoding. Participants completed a total of 5 blocks, each consisting of 18 trials. Within each block, participants performed 6 consecutive trials in one of the three virtual environments (baseline, vertical deformation, horizontal deformation) before switching to the next environment. The order of the environments was randomized within each block. This design resulted in a total of 90 test trials across all blocks (duration: M = 43.2 min, SD = 12.6 min).

Participants learned objects at 6 fixed target locations that were freely floating in volumetric space (Supplementary Fig. S1C). These locations and their distances to the nearest wall were evenly distributed across spatial dimensions, with the median coordinates close to the center of the environment and no significant difference in uniformity between spatial dimensions (Supplementary Fig. S1D-E). The minimum Euclidean distance to walls and between targets was 26 cm, resulting in a coverage of 5/8 of the space and smaller than larger inter-target distances (M = .747 m, SD = .585 m; Supplementary Fig. S1F). 3D models of everyday-life objects (camera, alarm clock, apple, measuring tape, wooden elephant, and teapot; Supplementary Fig. S1H) were taken from the *Poly Haven* asset library (www.polyhaven.com; published under the free license CC0). The mapping between objects and target locations was randomized across participants, and we observed differences in spatial memory error only for the target location, not for object identity (Supplementary Fig. S3A-B). Start locations were pseudo-randomly sampled so that the minimum distance between start and target location was 1 m and the entire environment was covered in each experiment phase. The order of trials was organized into sets of 6 trials, with each object-location sampled once in each set and no two consecutive trials containing the same object-locations. There was no time limit for completing trials and no virtual teleportation between trials.

### Debriefing

At the end of the experiment participants completed a short questionnaire on a computer screen. We asked participants about their deformation perception (whether the deformation was noticed, and if, the direction and magnitude of deformation), spatial perspective taking (egocentric vs allocentric vs mixed perspective), boundary-related strategies (fixed distance vs fixed ratio vs mixed strategy), as well as self-rated motion sickness, immersion, and motivation. Debriefing-related descriptions and results are summarized in Supplementary Fig. S6.

#### Data processing and analysis

Statistical analyses and visualization were performed in R 4.2.2 using the RStudio developer environment (version 2023.06.0). Unless otherwise noted, we used an alpha level of 5 % and two-sided tests. In case of multiple comparison testing, we applied the Bonferroni correction.

### Training and test performance

Baseline performance, i.e. spatial memory accuracy, in volumetric space was assessed by calculating the Euclidean distance in 3D space between the correct target location and the replacement locations. We first computed the error in spatial memory, averaged over target locations, for each repetition during the feedback phase. We then tested for differences across repetitions (pooled) and between walking and flying using a 1-way repeated measures ANOVA and a two-sample *t*-test, respectively. Successful encoding, resulting from training, was assessed by comparing the pooled spatial memory errors in the baseline test environment against the mean chance level via a 1-sample *t*-test. The chance level was determined by simulating 10,000 random responses for each target location and calculating the corresponding Euclidean error.

### Locomotion- and geometry-dependent anisotropy

To compare the precision of the horizontal and vertical encoding in volumetric space, we computed the median absolute deviation (MAD) for each target location within participant. This dispersion measure is robust to outliers, and does not require a reference to the correct value, making it suitable for deformed testing environments where responses cannot be compared to correct locations. The MAD reflects the consistency of spatial memory (analogous to the precision, or width of receptive fields of spatially-tuned cells). We compared the MAD for the z-axis (vertical dimension) to the average of the x and y axes (horizontal dimension) in two locomotion modes using a 2-way mixed ANOVA followed by pairwise post-hoc *t*-tests.

Next, we evaluated if the MAD is dependent on the distance between a target location and the environmental boundaries and if this relationship differs across vertical and horizontal dimensions as well as locomotion modes. Separately for the horizontal and the vertical dimension, and each participant, the distance of each object to the closest boundary was computed and we correlated these distance values with the MAD values. The individual Spearman correlation values were averaged for each locomotion group and tested against zero in a 1-sample *t*-test. Similarly, we tested for a relationship between the vertical MAD and the distance of a target to the participants’ body height (measured prior to the experiment; see Supplementary Table S1).

Subsequently, we tested how the response dispersion changes with deformations of the environment, that is, whether the change in MAD between baseline and the deformed environments is proportional to the actual deformation factor (i.e. 33%). To this end, we computed anisotropy indices for each participant and environment by taking the log-ratio of the MAD between the vertical and horizontal dimensions (positive or negative values indicated vertical or horizontal elongation, respectively, and a value of zero indicated isotropy). We then subtracted the baseline anisotropy from each deformed environment to quantify the deformation-induced change (i.e. rescaling). This deformation-induced MAD change was compared to the actual change in the aspect ratio of the environment (elongation by 1.33, compression by 0.67). Paired *t*-tests were performed for each deformed environment and locomotion mode separately.

### Navigational parameters

#### State occupancy

We divided the environment into 50 × 50 × 50 equally sized bins and calculated the number of visited bins in the baseline testing environment across trials and participants. Location visits were based on the position of a collector rigid body, since it is most comparable between both locomotion conditions.

#### Path length

The total path length between the starting location and replacement location in the baseline testing environment was computed as cumulative frame-wise 3D Euclidean distance and averaged for each participant.

#### Path dissimilarity

For each target location we computed the dissimilarity of movement paths between the starting and replacement location across trials via dynamic time warping (DTW, via the *dtw* function in *R*; Giorgino, 2009). DTW computes the similarity between two temporal sequences that may vary in speed or timing. It optimally aligns the sequences by stretching or compressing segments based on a warping function, thereby minimizing the cumulative distance between aligned points. First, the time series was downsampled from 40 Hz to 4 Hz. Then, we computed the DTW dissimilarity for each pair of trials for a particular target location before averaging the values for each participant.

#### Heading difference

We computed the circular average horizontal (azimuth) and vertical (elevation) orientation of the head at object placement during the feedback learning and test phase in the baseline environment and calculated its angular difference.

We tested for differences between the two locomotion modes in these parameters via two-sample *t*-tests. To evaluate if individual differences in navigational parameters are related to the processing of the vertical axis, we computed Spearman’s Rho correlation values between the vertical MAD of responses and each navigational parameter.

### Modeling

Participants can remember the location of objects using different geometric relations or features such as the absolute distance to the closest walls or the ratio of the distances to opposing walls. We tested which of several geometrical models best explains the object replacement data in the deformed environment (Hartley et al. 2004). In a deformed environment, locations that share features with the encoding environment are more likely to be chosen for object placement whereas locations with low feature similarity to the encoding environment are less likely to be chosen. For each location in the encoding environment, we can generate a prediction field with the shape of the testing environment that has this gradual likelihood. This was done for the following models:

#### Fixed distance

This model represents locations as feature vectors that contain a location’s perpendicular distance *d* to the closest wall on each environmental axis (TG: top-ground, FB: front-back, LR: left-right). In a 3D environment, this vector is of length 3:

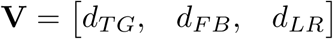

#### Fixed ratio

Locations are represented as three-dimensional vectors that encode the ratio of distances *r* to opposing walls on each environmental axis (e.g., *r*TG as the proportion of the object’s distance to the top wall to the distance between the top and ground wall).

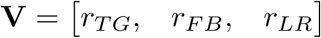

#### Boundary-proximity

The boundary-proximity model developed by Hartley and colleagues (2004) produces prediction fields similar to those derived from a population-model of boundary-vector cells (Hartley et al., 2000; Barry et al., 2006). Locations are encoded by their proximity to all walls (top, ground, front, back, left, right). Proximity is defined as 1/(*d* + *c*), with the distance *d* to a wall and the constant *c*. In a three-dimensional environment with six walls each location is represented by a six-dimensional vector:

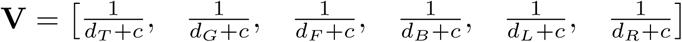

The boundary-proximity model behaves like a fixed distance model for objects close to the walls and like a fixed ratio model for objects at the center (see Hartley et al., 2004 and Fig. 3D-E). The c parameter controls the degree to which predictions align more closely with fixed distance or fixed ratio predictions. Like in Hartley and colleagues (2004) we set the constant *c* to the half of the cubic baseline environment’s axis size (1.5 m).

#### Ground-proximity

This model is a modification of the boundary-proximity model which reflects the direction of gravity in the 3D environment. Accordingly, we retained the distance to the floor and removed the distance to a ceiling in the model, resulting in a five-dimensional vector for each object-location:

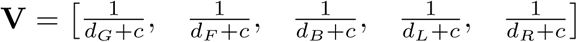

#### Model fit and comparison

We generated a prediction field (Fig. 3A; Supplementary Fig. S5) for each model to compute the likelihood of an object replacement in a given testing environment. First, the encoding environment was down-sampled to a 25 x 25 binned grid and the deformed environments accordingly (e.g., 33 x 25 binned grid in a horizontally stretched environment). Then, we computed the model-specific feature vectors for each location in the encoding (cube) as well as the deformed environments (rectangular cuboids). The *L^2^* norm between two vectors, one from the encoding and one from the testing environment, was computed and the repetition of this process for all cross-environment combinations yielded a representational dissimilarity matrix *D*xyz. This matrix was then converted into a similarity matrix by:

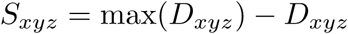

*S*xyz was turned into a probability matrix *P*xzy by applying a softmax function:

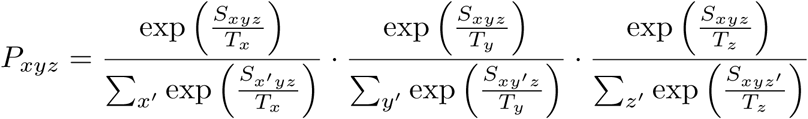

where *T* are the dimension-specific temperature parameters that control the sharpness of the probability distribution (low values approximating peak predictions and high values uniformity). The three temperature parameters were first fit to the object replacements in the baseline test environment, for each model and participant separately, using the Limited-memory BFGS algorithm (Liu & Nocedal, 1989) via the minimize function of the *scipy* Python package.

The fitted models were subsequently applied to the deformed environments to receive a log-likelihood (LL) estimate for each participant and model. The LL estimates were compared between the models in repeated-measures ANOVAs and subsequent post-hoc *t*-tests (with Bonferroni-based family-wise error correction). The modeling analyses and visualizations were run by custom Python (3.9) scripts.

## Acknowledgments

We would like to thank Max Schulz and Paula Schneider for helping with the recruitment of participants and data collection. We also thank Kiran Varanasi for helpful input on setting up the MoCap-featured VR lab. This work was supported by the Max Planck Society. C.F.D. is supported by the Max Planck Society, the Kavli Foundation, the Jebsen Foundation and Helse Midt Norge.

## Author contributions

*VR* and *TS* contributed equally to this work.

*VR:* Project idea, Conceptualization, Data acquisition, Formal analysis, Visualization, Software, Task programming, Writing - original draft, Writing - review & editing

*TS:* Conceptualization, Formal analysis, Modeling analysis, Visualization, Software, Writing - original draft, Writing - review & editing

*LK:* Task programming, Software, Writing - review & editing

*MK:* Conceptualization, Supervision, Writing - review & editing

*CD:* Conceptualization, Supervision, Resources, Writing - review & editing

## Data availability

Data to reproduce the statistical analyses reported in this paper will be made available upon publication via the Open Science Framework.

## Code availability

The analysis code will be available upon publication on GitHub.

## Supplemental information

**Figure S1.**
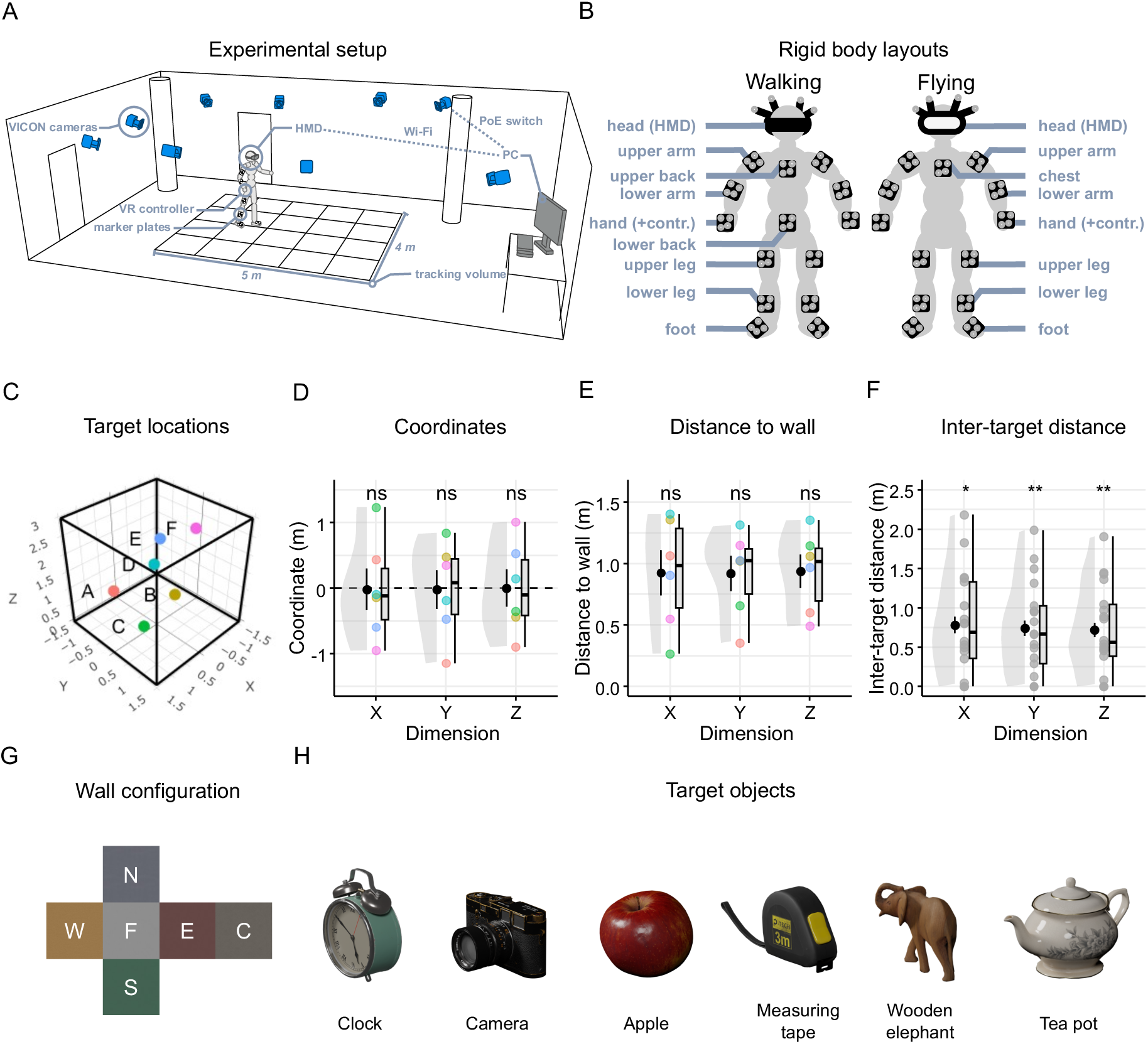
Experimental setup and predefined object-locations. **A:** Schematic of the Motion-Capture (MoCap)-featured VR lab and used equipment. **B:** Rigid body layouts for participants of the flying (13 rigid bodies) and walking (14 rigid bodies) groups, illustrating the placement of markers on different parts of the body. Note that the flying participants did not wear a marker on their lower back because they were sitting in a rotating chair whose position we tracked instead (not shown here). **C:** Fixed set of six target locations (colored and labeled dots) within the cubic baseline environment. **D-F:** We assessed whether the target-related values (coordinates, distance-to-wall, inter-target distance) within each dimension were uniformly distributed using the Kolmogorov-Smirnov (KS) test. To compare uniformity across the dimensions, we performed pairwise permutation tests. For each comparison, we calculated the absolute difference between the KS statistics of the two dimensions. We then generated 1,000 permuted datasets by randomly shuffling the distances between dimensions and recalculated the KS statistic differences. **D:** Coordinate distribution of target locations (colored dots) for each spatial dimension. Coordinates of any dimension did not significantly deviate from uniformity (*KS*-test for uniformity; X-dimension: *D* = .226, *P*_adj_ = 1, Bonferroni-corrected for 3 comparisons; Y-dimension: *D* = .167, *P*_adj_ = 1; Z-dimension: *D* = .168, *P*_adj_ = 1) with no differences in the magnitude of uniformity across dimensions (Permutation-based comparison between pairs of dimensions: Mean Δ*D* = .04 m, all *P*_adj_’s = 1, Bonferroni-corrected for 3 comparisons). **E:** Distribution of distances between targets (colored dots) and nearest wall for each spatial dimension. Distances of any dimension did not significantly deviate from uniformity (X-dimension: *D* = .293, *P*_adj_ = 1, Bonferroni-corrected for 3 comparisons; Y-dimension: *D* = .333, *P*_adj_ = 1; Z-dimension: *D* = .284, *P*_adj_ = 1) with no differences in the magnitude of uniformity across dimensions (Mean Δ*D* = .096 m, all *P*_adj_’s < .744, Bonferroni-corrected for 3 comparisons). **F.** Distribution of distances between target locations for each spatial dimension. Distances were non-uniformly distributed (X-dimension: *D* = .276, *P*_adj_ = .025, Bonferroni-corrected for 3 comparisons; Y-dimension: *D* = .308, *P*_adj_ < .01; Z-dimension: *D* = .3, *P_adj_* < .01) with more small than large inter-target locations but no differences between dimensions (Mean Δ*D* = .017 m, all *P*_adj_’s = 1, Bonferroni-corrected for 3 comparisons). Violin plots depict the density distribution, boxplots the median and quartiles, black dots with error bars the means ± SEM, and colored dots individual data points per condition. * *P* < .05, ** *P* < .01. **G:** Example baseline wall configuration; F = Floor, C = Ceiling, N = North, S = South, E = East, W = West. **H:** 3D models of target objects used in the experiment.

**Figure S2.**
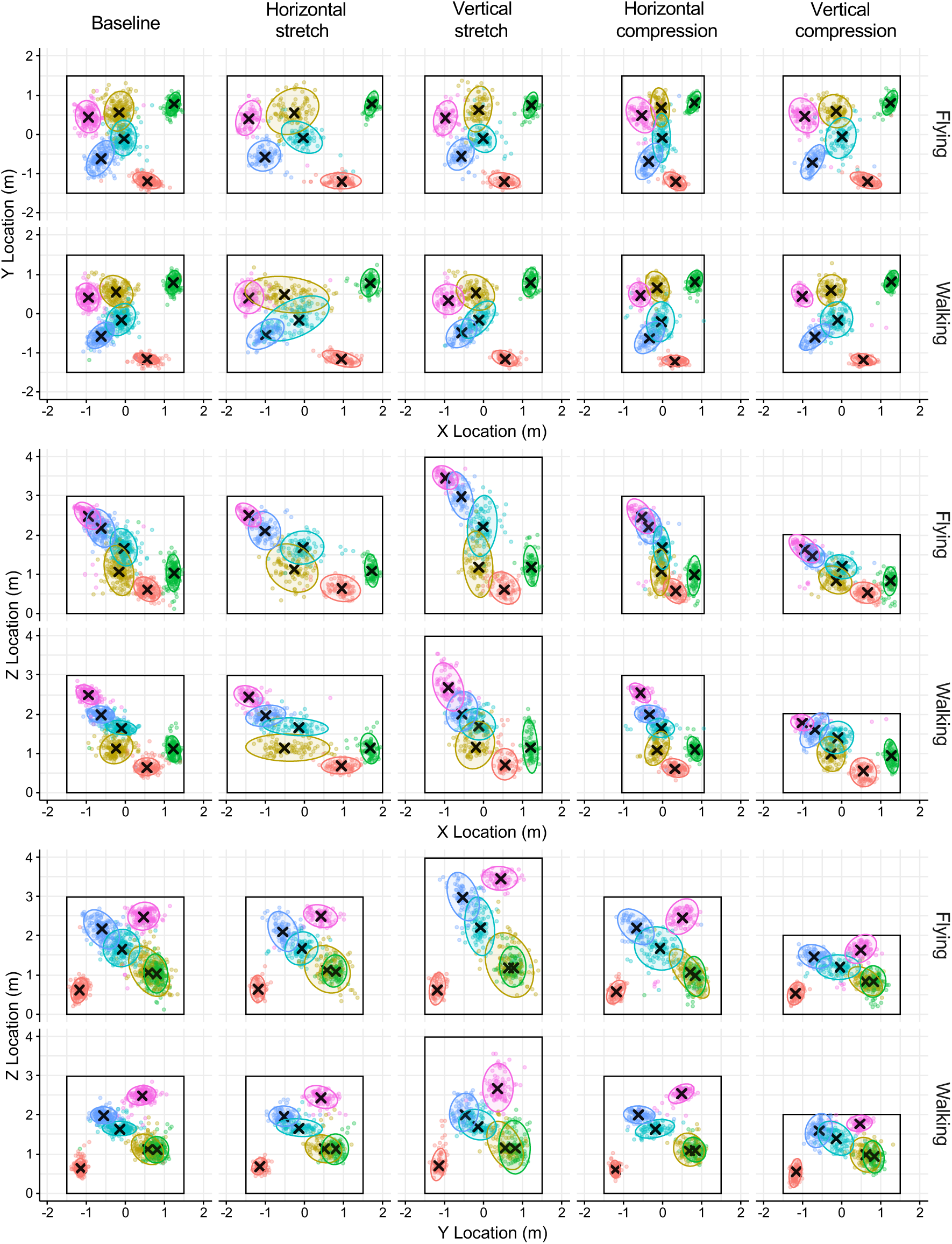
2D projections of object replacements at test. Object replacements (colored dots) in the baseline environment (first column) and deformed environments (columns 2-5) of the flying and walking groups (first and second row of each subplot, respectively). Each dot represents one participant’s single response, with 5 replacements for each target location. Colored ellipsoids based on multivariate *t-*distributions, each covering 95% of the data, were fitted to each target’s median response (superimposed crosses). Data is separately plotted for each 2D projected view: Top view (XY; row 1-2), front view (XZ; row 3-4), side view (YZ; row 5-6).

**Figure S3.**
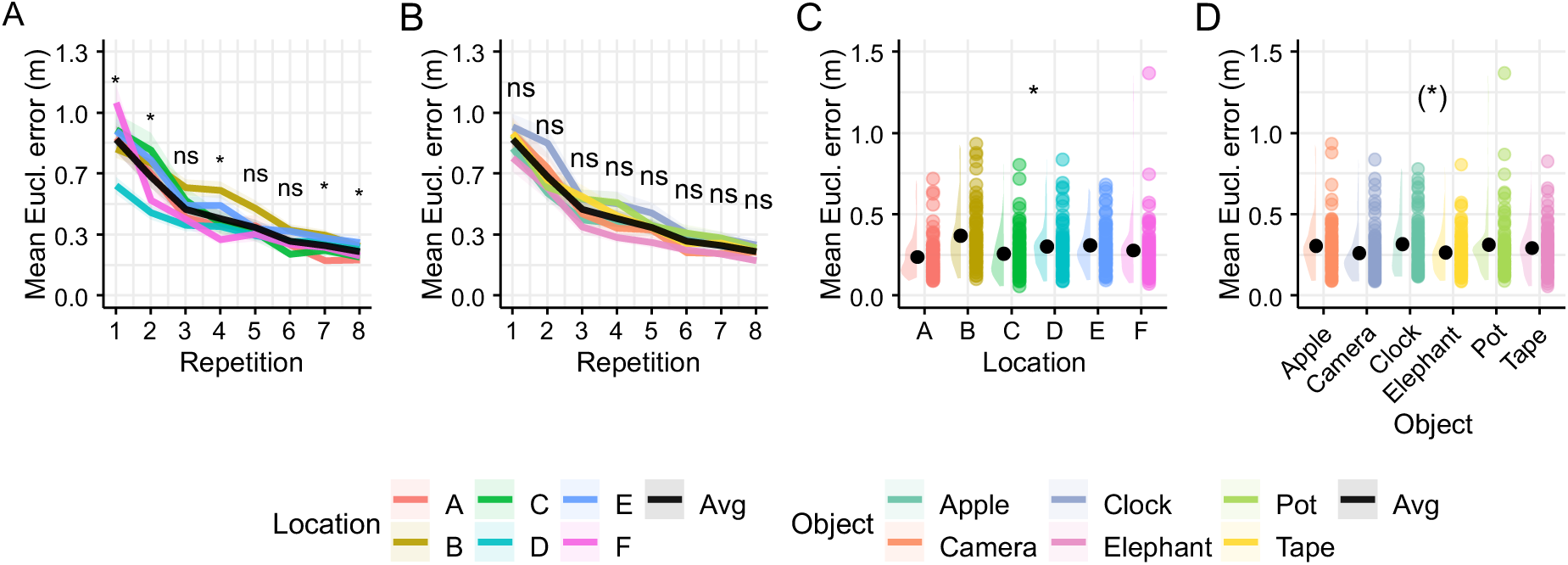
Location- and object-specific spatial memory performance at baseline. **A:** Training accuracy (mean Euclidean error) of each target location as a function of repetition. Errors significantly differed between target locations at all repetitions, except for 3, 5 and 6 (1-way repeated measures ANOVAs: all *P*’s < .05, Bonferroni-corrected for 8 comparisons). **B:** Training accuracy (mean Euclidean error) of each object (e.g. apple, clock) as a function of repetition. Errors did not differ between objects at all repetitions (1-way repeated measures ANOVAs: all *P*’s > .10, Bonferroni-corrected for 8 comparisons). **C:** Test accuracy for each target location. Errors were significantly different across target locations (1-way repeated measures ANOVA: *F*_4.31,327.23_ = 10.356, *P* <.001, 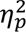 = .12). **D:** Test accuracy for each object. Although errors differed weakly between objects (*F*_4.37,332.47_ = 2.544, *P* = .04, 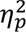 = .032), none of the post-hoc pairwise comparisons survived Bonferroni-correction for multiple comparisons (all *P’*s > .2). This, along with the small effect size, suggests that differences in errors across objects are not statistically robust. Color-coded lines depict means ± SEM. Violin plots depict the density distribution, black dots with error bars the means ± SEM, and colored dots individual data points per condition. * *P* < .05.

**Figure S4.**
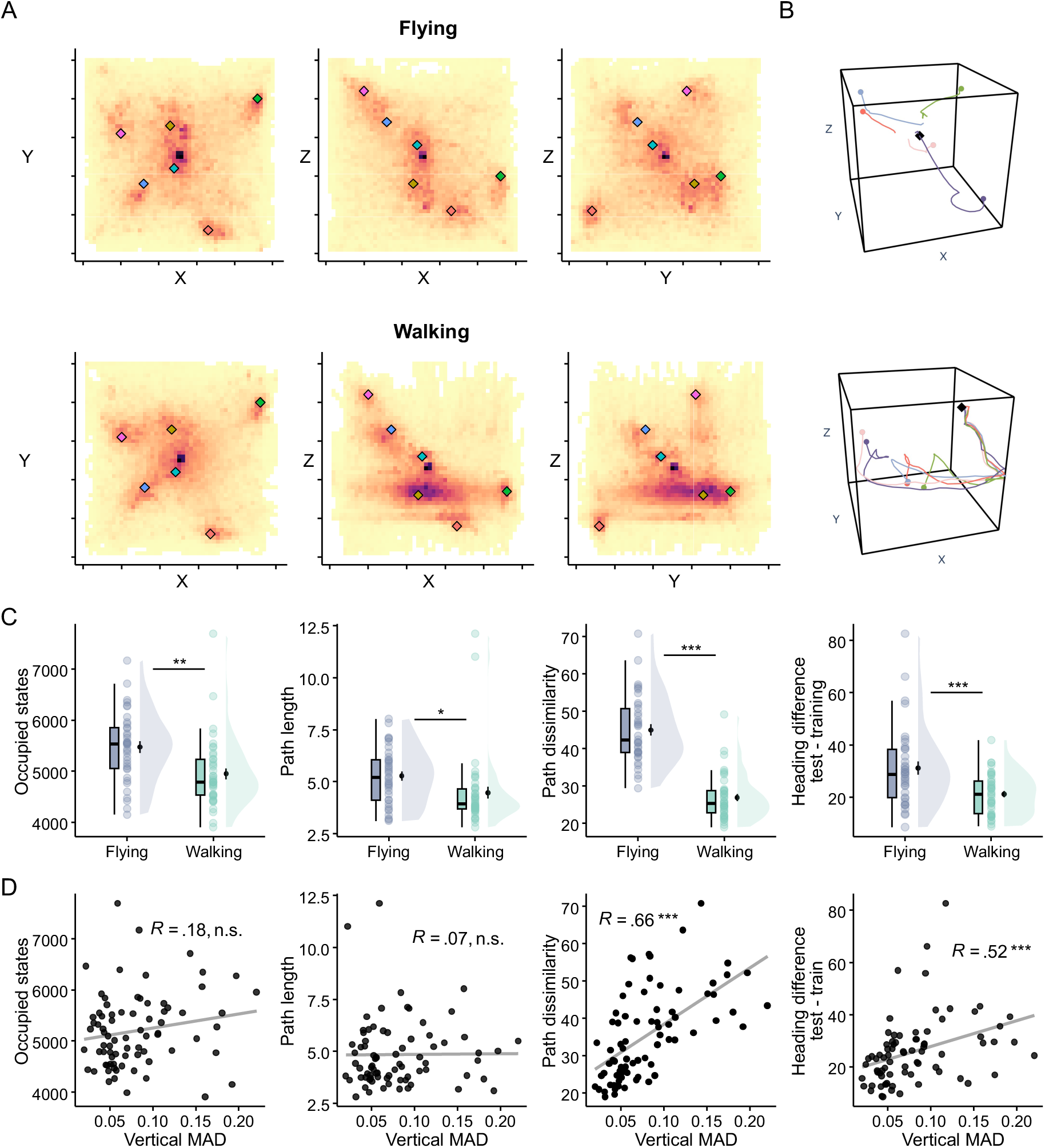
Navigation behavior differs between locomotion modes. **A:** Location-occupancy of the collector in the baseline environment (downsampled to 50 bins), normalized within and summed across participants and projected onto 2D planes, with darker shades reflecting more location visits. Colored diamonds depict the target locations. **B:** Example trajectories of two participants (top: flying; bottom: walking) in the baseline environment engaging distinct navigational strategies. Colors refer to different trials, circles to the starting location and the black diamond to the target location. **C:** Locomotion-dependent differences in state occupancy (left; *t*_74.5_ = 3.38, *P* = .001, *d* = .771), path length (center-left; *t*_71.8_ = -2.24, *P* = .028, *d* = -.508), path dissimilarity (center-right; *t*_61.7_ = 10.1, *P* < .001, *d* = 2.32) and test vs. training heading difference (right; *t*_52.4_ = 3.57, *P* < .001, *d* = .823). **D:** Correlations (Spearman’s Rho) between navigational variables from C with vertical median absolute deviation (State occupancy: Spearman’s *Rho* = .18, *S* = 62696, *P* = .126; Path length: Spearman’s *Rho* = .07, *S* = 70874, *P* = .554; Path dissimilarity: Spearman’s *Rho* = .66, *S* = 25672, *P* < .001; Heading difference: Spearman’s *Rho* = .52, *S* = 36594, *P* < .001; all other *P* > .126). * *P* < .05, ** *P* < .01, *** *P* < .001.

**Figure S5.**
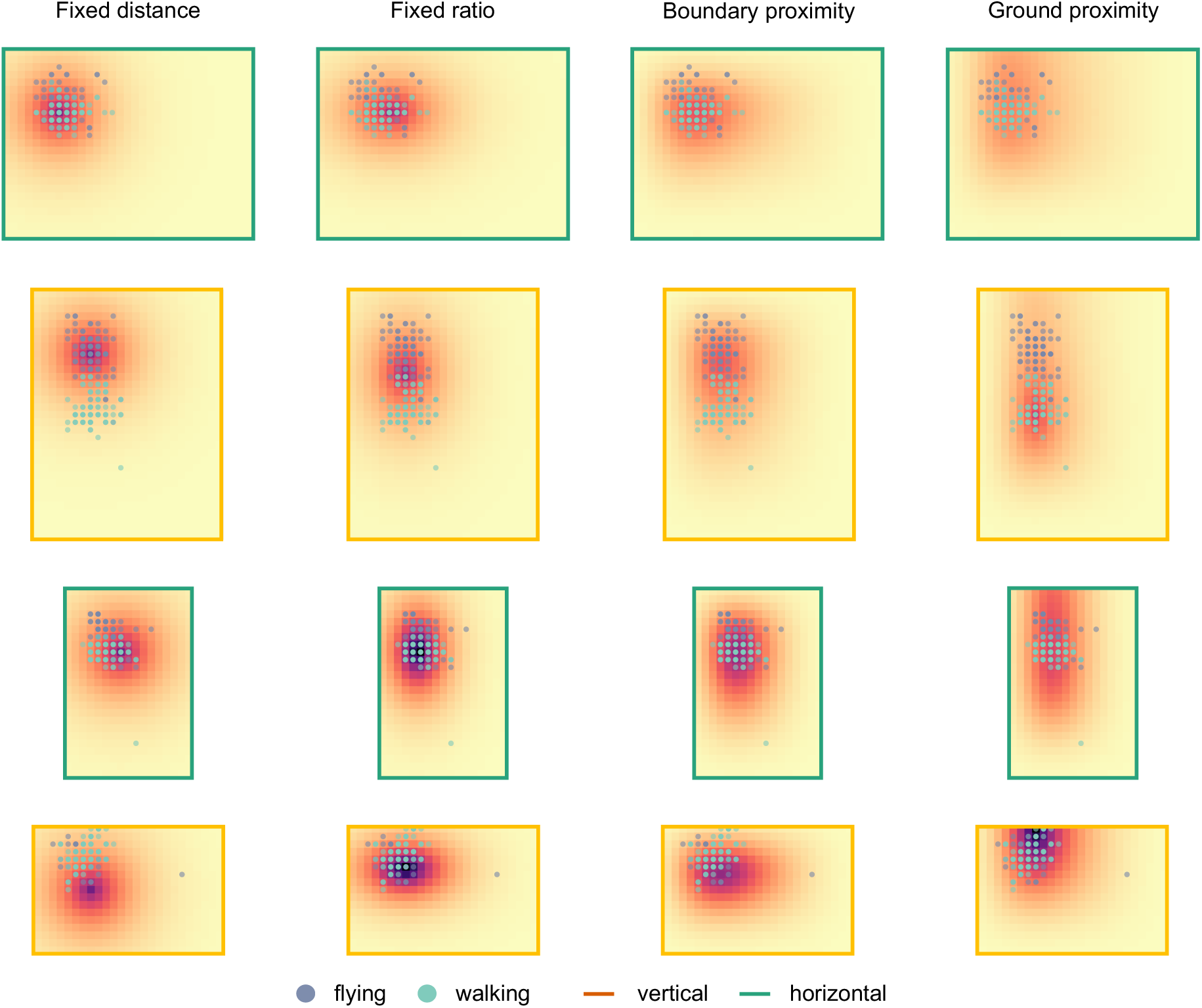
Model prediction fields. Heatmaps represent the (unfitted) prediction fields derived from various geometric models (column 1: *Fixed distance*, 2: *Fixed ratio*, 3: *Boundary-proximity*, 4: *Ground-proximity*) under different environmental deformations (rows 1-2: stretch, 3-4: compression) for one example target location (see Fig. 1E, object E). Darker colors indicate areas of higher replacement probability. Superimposed dots depict binned object replacements of walking (cyan) and flying (grayish blue) participants. The color of the frames reflects the type of environmental deformation: vertical (orange) versus horizontal (green). Prediction fields of each model were fitted to object replacements of walking and flying participants (see Methods section under “Modeling”).

**Figure S6.**
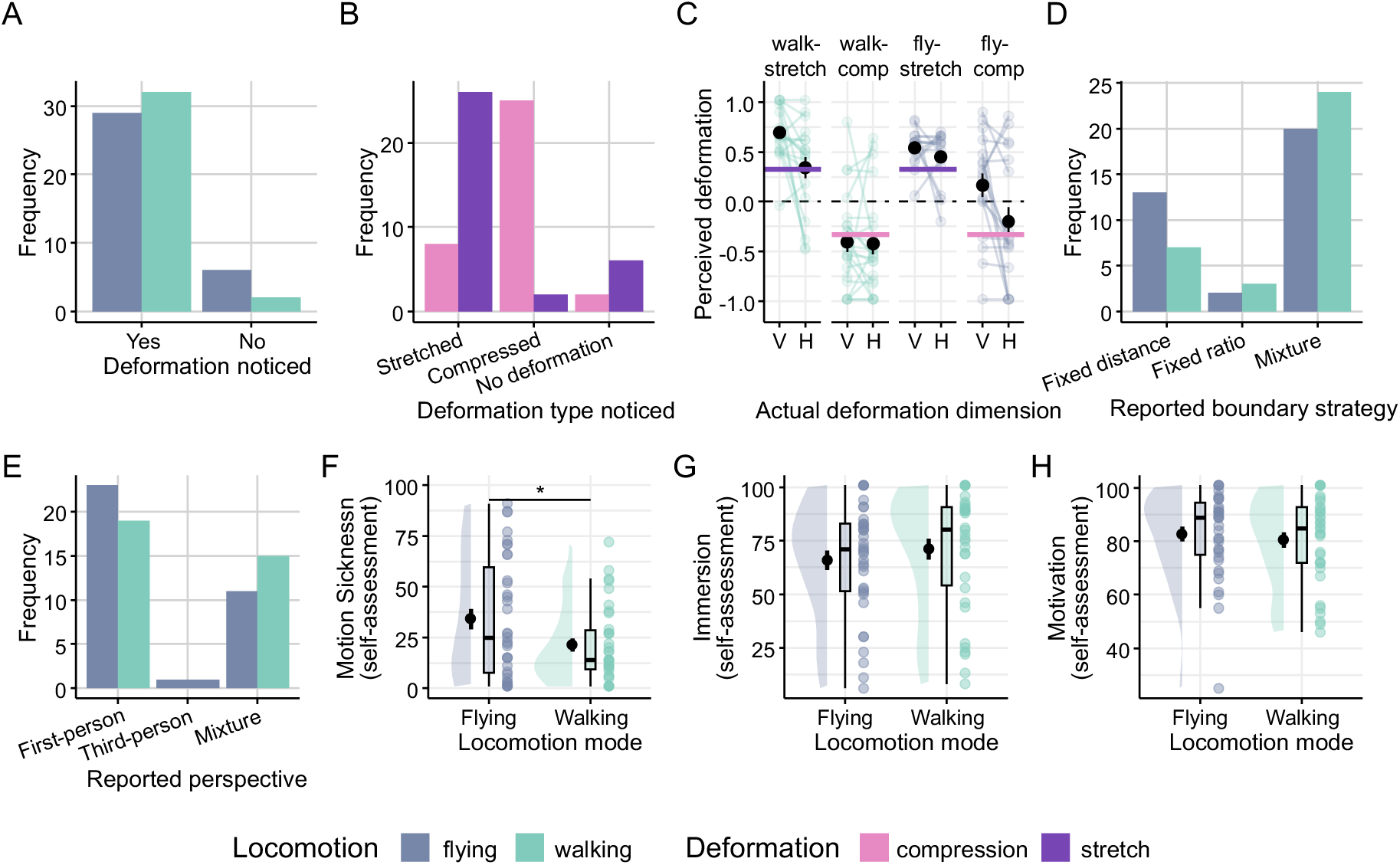
Debriefing summary. **A:** Number of participants who noticed the deformation of the virtual environment during the test phase. The majority (89.6%) perceived the deformation. Question: “*Did you notice that the virtual environment changed during the test phase?”*; Options: “*Yes*” / “*No*” **B:** Number of participants who correctly identified the type of deformation [‘Stretch’, ‘Compression’, or ‘None’]. 76.6% of participants who perceived the deformation correctly identified the deformation type. Question: “*In which way did the direction change?*”; Options: “*The size of the environment has been increased*”, “*The size of the environment has been decreased*”. **C:** Group-specific perception of the deformation magnitude (negative values = compression; positive values = stretch; 0 = no change) in the vertical (V) and horizontal (H) dimensions, compared to the actual deformation magnitude (horizontal lines). Most participants tended to correctly estimate horizontal deformations, while those in the walking condition tended to overestimate vertical stretch and those in the flying condition tended to overestimate vertical compression. Question: “*How much did the size change?*”; Options: Bidirectional continuous slider on a visual scale for vertical and horizontal dimension, separately (center = no change relative to baseline, minimum = half the size of baseline, maximum = 2 times baseline). Black dots depict mean ± SEM with error bars, color-coded dots reflect individual data points per condition. **D:** Number of participants who reported using different boundary-related strategies to remember object-locations. The majority (63.08%) of participants reported using a mixture between ‘fixed distance’ and ‘fixed ratio’ strategy Question: “*How did you use the walls of the virtual box to remember the location of the objects?”;* Options: “*I remembered the edge to the nearest walls”* [‘Fixed distance’] / “*I remembered the ratio of opposite walls”* [‘Fixed ratio’] / “*A mixture of the two options*” [‘Mixture’]. **E:** Number of participants who reported using different spatial perspective-taking strategies. 57.14% of participants reported using an egocentric perspective, 1.43% using an allocentric strategy and 41.43% using a mixture strategy, with comparable proportions between locomotion modes. Question: “*What strategy did you use to remember the object-locations?”;* Options: “*I tried to remember a snapshot (‘first-person perspective’)”* [‘Egocentric’] / “*I tried to imagine a mental ‘map’ of the virtual box in order to locate objects on the ‘map’ (‘third-person perspective’)”* [‘Allocentric’] / “*A mixture of the first two options*” [‘Mixture’] **F:** Self-reported motion sickness levels, with flying participants reporting significantly higher motion sickness compared to walking participants (2-sample *t*-test: *t*_58.8_ = 2.27, *P* = .027, *d* = .523). Question: “*Did you feel dizzy or nauseous while performing the task”;* Options: Continues slider on a visual scale from “not at all” to “very much”. Violin plots depict the density distribution, boxplots the median and quartiles, mean ± SEM as black dots with error bars, as well as individual data points (dots) per condition. * *P* < .05. **G:** Self-reported immersion levels, with no significant differences between the flying and walking groups (2-sample *t*-test: *t*_75_ = -.572, *P* = .569, *d* = -.130). Question: “*How much did you feel that you were in the ‘here and now’ while performing the task?”;* Options: Continues slider on a visual scale from “not at all” to “very much”. **H:** Self-reported motivation levels, with no significant differences between the flying and walking groups (2-sample *t*-test: *t*_68.4_ = -.482, *P* = .631, *d* = -.110). Question: “*How high would you rate your motivation during the experiment?”;* Options: Continues slider on a visual scale from “not at all” to “very much”.

**Table S1.** Sample characteristics.

| Variable | Group 1 (n=20) | Group 2 (n=18) | Group 3 (n=20) | Group 4 (n=19) | Full sample (N=77) | P-value |
| --- | --- | --- | --- | --- | --- | --- |
| **Gender** |  |  |  |  |  |  |
| Female | 9 (45%) | 10 (55.6%) | 12 (60%) | 9 (47.4%) | 40 (51.9%) | .780 |
| Male | 11 (55%) | 8 (44.4%) | 7 (35%) | 10 (52.6%) | 36 (46.8%) | .611 |
| Non-binary | 0 (0%) | 0 (0%) | 1 (5%) | 0 (0%) | 1 (1.3%) | 1 |
| **Age** |  |  |  |  |  | .408 |
| 18-23 years | 6 (30%) | 9 (50%) | 3 (15%) | 4 (21.1%) | 22 (28.6%) | .106 |
| 24-29 years | 10 (50%) | 5 (27.8%) | 12 (60%) | 12 (63.2%) | 39 (50.6%) | .132 |
| 30-35 years | 4 (20%) | 4 (22.2%) | 5 (25%) | 3 (15.8%) | 16 (20.8%) | .945 |
| **Education** |  |  |  |  |  |  |
| < High school | 1 (5%) | 4 (22.2%) | 0 (0%) | 1 (5.3%) | 6 (7.8%) | .08 |
| High school | 7 (35%) | 7 (38.9%) | 11 (55%) | 10 (52.6%) | 35 (45.5%) | .546 |
| Bachelor's | 10 (50%) | 6 (33.3%) | 3 (15%) | 4 (21.1%) | 23 (29.9%) | .09 |
| Master's | 2 (10%) | 1 (5.56%) | 5 (25%) | 4 (21.1%) | 12 (15.6%) | .322 |
| **Body measures** |  |  |  |  |  |  |
| Weight (kg) | 66.2 ± 9.44<br>(45 - 89) | 70.2 ± 10.1<br>(58 - 85) | 71.3 ± 12.4<br>(52 - 94) | 73.9 ± 12.6<br>(51 - 100) | 70.36 ± 11.38<br>(45 - 100) | .206 |
| Height (m) | 1.75 ± .076<br>(1.65 - 1.87) | 1.77 ± .05<br>(1.69 - 1.85) | 1.73 ± .061<br>(1.63 - 1.84) | 1.77 ± .07<br>(1.65 - 1.88) | 1.76 ± .07<br>(1.63 - 1.88) | .196 |
| Arm span (m) | 1.72 ± .099<br>(1.58 - 1.92) | 1.75 ± .07<br>(1.6 - 1.86) | 1.72 ± .08<br>(1.6 - 1.85) | 1.75 ± .09<br>(1.57 - 1.9) | 1.74 ± .083<br>(1.57 - 1.92) | .425 |
| Inseam (m) | .794 ± .046<br>(.73 - .88) | .817 ± .06<br>(.74 - .98) | .785 ± .03<br>(.73 - .83) | .795 ± .04<br>(.7 - .85) | .797 ± .045<br>(.7 - .98) | .15 |
| Hip width (m) | .368 ± .028<br>(.34 - .46) | .374 ± .022<br>(.33 - .4) | .376 ± .021<br>(.33 - .41) | .371 ± .025<br>(.34 - .42) | .372 ± .024<br>(.33 - .46) | .668 |

*Note.* Gender, age, and education: Values depict frequency (percentage) for each group and pooled across groups; *P*-values resulted from Fisher's exact test (two-sided) assessing associations with group membership. Body measures: Values depict mean ± SD (min-max) for each group and pooled across groups. *P*-values resulted from 1-way ANOVAs, each with group membership as a between-subject factor. Group 1 = Walking-Stretch; Group 2 = Flying-Stretch; Group 3 = Walking-Compression; Group 4 = Flying-Compression.

**Table S2.** Environment characteristics.

| Attribute | Environment A | Environment B | Environment C | Environment D | Environment E |
| --- | --- | --- | --- | --- | --- |
| Geometric shape | Cube | Cuboid | Cuboid | Cuboid | Cuboid |
| Deformation dimension | - | Horizontal (X) | Vertical (Z) | Horizontal (X) | Vertical (Z) |
| Deformation type | - | Stretch | Stretch | Compression | Compression |
| Group | All | 1, 2 | 1, 2 | 3, 4 | 3, 4 |
| Phase | Training & Test | Test | Test | Test | Test |
| X/length (m) | 3 | 3.99 | 3 | 2.01 | 3 |
| Y/width (m) | 3 | 3 | 3 | 3 | 3 |
| Z/height (m) | 3 | 3 | 3.99 | 3 | 2.01 |
| Re-scaling (m) | 0 | .99 (33%) | .99 (33%) | .99 (33%) | .99 (33%) |

